# Cytokine mRNA Delivery and Local Immunomodulation in the Placenta using Lipid Nanoparticles

**DOI:** 10.1101/2025.02.07.637086

**Authors:** Samuel I. Hofbauer, Luisa A. Fink, Rachel E. Young, Tara Vijayakumar, Katherine M. Nelson, Nia Bellopede, Mohamad-Gabriel Alameh, Drew Weissman, Jason P. Gleghorn, Rachel S. Riley

## Abstract

During pregnancy, the maternal immune system adapts to balance tolerance of the semi-allogenic fetus while protecting the fetus from pathogens. Dysregulated immune activity at the maternal-fetal interface contributes to pregnancy complications, such as recurrent pregnancy loss and preeclampsia. Compared to healthy placentas, preeclamptic placentas exhibit increased pro-inflammatory signaling, including a predominance of inflammatory macrophages, leading to impaired tissue remodeling and restricted blood flow. However, the precise mechanisms driving this immune imbalance remain poorly understood, in part due to the lack of tools to probe individual pathways. Here, we use lipid nanoparticles (LNPs) to deliver cytokine-encoded mRNA to placental cells, called trophoblasts, enabling local immunomodulation. LNP-mediated delivery of IL-4 and IL-13 mRNA induced cytokine secretion by trophoblasts, leading to polarization of primary human monocytes toward anti-inflammatory phenotypes. Notably, lowering the mRNA dose increased expression of alternatively-activated macrophage markers, revealing an inverse relationship between dose and polarization status. Intravenous injection of LNPs in pregnant mice achieved placental secretion of IL-4 and IL-13 with minimal changes to pro-inflammatory cytokines in the serum. These findings establish LNPs as a tool for local immunomodulation in the placenta, offering a strategy to study and treat immune dysfunction in pregnancy and in other inflammatory conditions.

## Introduction

Nanoparticle-based therapeutics, and in particular lipid nanoparticles (LNPs), have recently emerged as tools to treat diseases of pregnancy. These platforms have been designed to either passively or actively accumulate in the placenta to deliver therapeutic nucleic acids and reverse the maternal symptoms of preeclampsia.^1–11^ In preeclampsia, pregnant patients present with new onset hypertension and proteinuria, and severe disease can lead to other end-organ damage, seizures, and possible death for the pregnant person and fetus. While the precise pathophysiology of preeclampsia is still under investigation, data supports that abnormal placenta development early in pregnancy leads to preeclampsia symptoms at later stages of gestation (>20 weeks).^12–14^ Thus, by the time symptoms arise, especially in severe cases, the underlying placenta damage is already advanced, leaving limited opportunity for gene therapy to effectively reverse the pathological processes. To develop therapeutics for preeclampsia, we first need to develop tools to understand the biological processes underlying healthy and pathologic placentation. Here, we aim to demonstrate the capability of LNPs to deliver cytokine mRNA to trophoblasts to serve as tools for local immunomodulation in the placenta.

The placenta plays several critical roles during pregnancy including nutrient and oxygen exchange, waste removal, and regulating immune activity. Immune activity within the uterine microenvironment and placenta is a delicate balance to support tolerance to the semi-allogeneic fetus, protection against infections, and placental invasion into the myometrium.^15^ Dysregulated immune activity is associated with several pregnancy complications including preeclampsia, intrauterine growth restriction (IUGR), and recurrent pregnancy loss.^16–21^ For example, in preeclampsia, there is a predominance of classically-activated macrophages,^21,22^ higher Th1:Th2 ratio and activity,^23,24^ and higher levels of natural killer (NK) 1 cells^25,26^ compared to healthy pregnancies. Thus, maintaining a healthy balance of pro- and anti-inflammatory signals in the placenta is critical to support placenta and pregnancy establishment. However, the underlying biological and immunological processes that lead to healthy or abnormal placenta development are still under investigation and are often controversial.^21,27,28^

Our primary focus in this work is on macrophage activity due to their central role in placental health and disease, as well as their roles in tissue repair and modulating immune responses.^29^ Macrophages exhibit a high degree of plasticity to regulate inflammation by adopting either pro-inflammatory or anti-inflammatory phenotypes depending on the signals they receive from their environment.^20,29^ Outside of pregnancy, macrophage polarization is being investigated in cancers, autoimmune disorders, wound healing, and others applications.^30–36^ However, understanding the role of macrophages during healthy and pathologic pregnancies is in early stages. In healthy pregnancies, macrophages support tissue remodeling, placental vascularization, and immune tolerance at the maternal-fetal interface.^20,37^ Conversely, in pregnancy-related disorders, such as preeclampsia, macrophage function favors the pro-inflammatory phenotypes, contributing to poor placental development and adverse pregnancy outcomes.^38–40^ While data clearly suggests a predominance of pro-inflammatory macrophages in the preeclamptic placenta, the mechanisms that underlie this immune dysregulation, and how this impacts placental development, remain poorly understood.

We aimed to establish LNPs as tools to study the underlying immunological mechanisms and to explore the therapeutic potential of immunomodulation in the placenta, particularly through macrophage polarization. It is well established that stimulation with IL-4 and IL-13 recombinant peptides polarize macrophages to an alternatively activated, anti-inflammatory state.^41–44^ After IL-4/IL-13 stimulation, alternatively activated macrophages secrete anti-inflammtory factors such as IL-10.^45^ IL-10 is critical for promoting maternal immune tolerance and regulating immune cell function to support tissue remodeling and trophoblast invasion.^46^ Of note, studies have found reduced IL-4 and IL-10 levels in serum and placenta tissue during preeclamptic pregnancies compared to normotensive pregnancies.^47–49^ Further, peripheral blood mononuclear cells (PBMCs) collected from preeclamptic patients secrete lower levels of IL-10 compared to those from healthy patients, indicating a higher Th1 response in preeclampsia.^50^ Thus, modulating the IL-4/IL-13 axis represents a means to both control macrophage phenotype and elicit anti-inflammatory activity in tissues by modulating IL-10 levels.

Here, we ecapsulated IL-4 and IL-13 mRNA inside LNPs to modulate this immunological axis in the placenta, establishing their use as tools to study the immunological basis of diseases of pregnancy. Prior studies have developed LNPs for delivering mRNA encoding angiogenesis-promoting factors, such as placental growth factor (PlGF) or VEGF, as potential therapeutic agents for preeclampsia.^1,4,8^ While these prior studies have focused on reversing preeclampsia symptoms, the current study represents the first evaluation of LNPs for cytokine secretion and immunomodulation in pregnancy. We used an LNP platform previously designed in our lab to deliver mRNA to trophoblasts *in vitro* and *in vivo*.^1^ LNP-induced secretion of IL-4 and IL-13 *in vitro* yielded alternatively-activated macrophage polarization of primary human monocytes, with an inverse relationship between administered mRNA dose and extent of polarization. Further, we demonstrate that intravenous delivery of LNPs in healthy pregnant mice achieves placental delivery and local cytokine secretion. Together, these findings lay the foundation for LNPs to modulate placental immune activity, highlighting their potential as tools to both study and treat immunologically-driven diseases of pregnancy.

## Results

### LNP Formulation and Characterization

In this work, we used an LNP formulation developed in our lab through a Design of Experiments approach that achieves preferential delivery to mouse placentas (Fig. 1a).^1^ This LNP is comprised of the C12-200 ionizable lipid (35%), DOPE (10%), cholesterol (53.5%), and DMPE-PEG (1.5%). We used this LNP formulation to deliver cytokine mRNA to the placenta for local immunomodulation. The ionizable lipids within LNPs are pH-responsive to become positively charged in acidic environments. This feature is important for the encapsulation of the negatively charged mRNA molecules and endosomal escape inside cells.^51^ We used the C12-200 ionizable lipid because it is well established for mRNA delivery pre-clinically by us and others.^1,52,53^ The mRNA encapsulated in the LNPs encoded human IL-4 (hIL-4), human IL-13 (hIL-13), murine IL-4 (mIL-4), murine IL-13 (mIL-13), or firefly luciferase (Luc), each of which were used in the experiments described herein. We characterized all LNPs used in this work for size by dynamic light scattering (DLS), which resulted in sizes ranging from 80.82-136.9 nm and low polydispersity indices <0.27 (Fig. 1b,c). Average mRNA encapsulation ranged from 44.19-82.26% (Fig. 1d), as assessed by Ribogreen assays.

**Figure 1.**
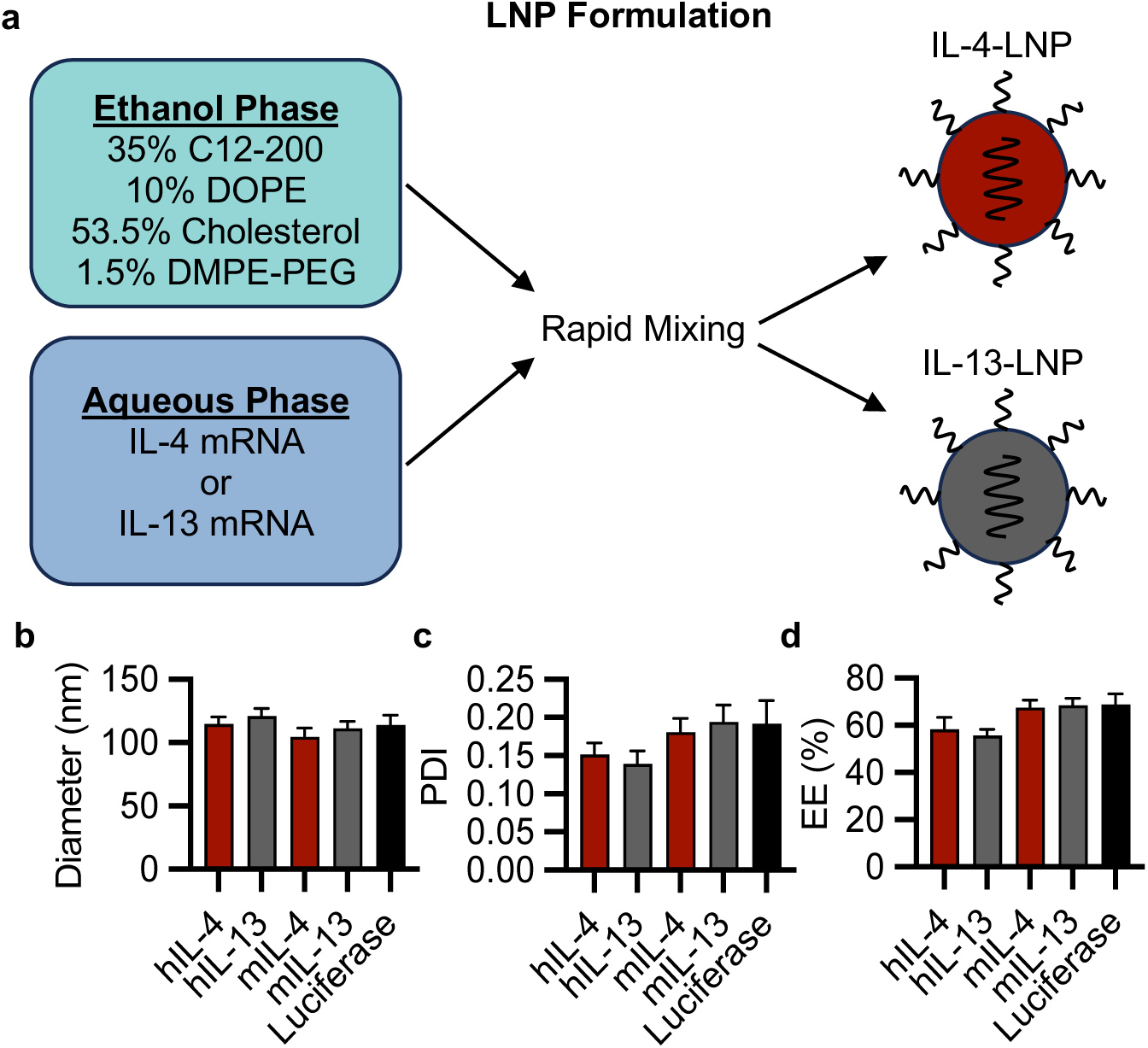
LNP formulation and characterization. a) Schematic depicting LNP formulation using our top platform for placenta delivery. LNP characterization including (b) mean hydrodynamic diameter, (c) polydispersity indices, and (d) encapsulation efficiencies of LNPs as assessed by Ribogreen assays.

### LNPs Enabled IL-4 and IL-13 mRNA Delivery to Trophoblasts *In Vitro*

We investigated LNP-mediated delivery of hIL-4 (hIL-4-LNP) and hIL-13 (hIL-13-LNP) to trophoblasts to induce ectopic cytokine secretion *in vitro*. Delivery in three trophoblast cell lines was evaluated including the b30 subclone of BeWo cells (termed BeWo herein), JAR cells, and HTR8/SVneo cells (HTR8 herein). BeWo and JAR cells are 3^rd^-trimester human choriocarcinoma cells,^54,55^ and HTR8 cells are a 1^st^-trimester trophoblast cell line.^55^ All three cell lines were used to assess the potential for LNPs to deliver mRNA to cells that represent different stages of gestation. In these experiments, cells were treated with hIL-4-LNPs or hIL-13-LNPs, and reported doses represent the total amount of mRNA, with half of the dose being IL-4-LNPs and half being IL-13-LNPs. BeWos and JARs were treated with LNPs at 0-600 ng mRNA/well, and HTR8s were treated with 0-1200 ng mRNA/well in 96-well plates (0-12 ng/μL mRNA) (Fig. 2). All three cell lines secreted cytokine in a dose-dependent manner. BeWos and HTR8s secreted higher levels of both cytokines compared to JARs. At the highest dose administered (600 ng total mRNA), BeWos secreted up to ∼295 ng of IL-4 and ∼102 ng of IL-13. Similarly, HTR8s treated with 1200 ng mRNA secreted up to 813.3 ng of IL-4 and ∼418 ng of IL-13. Lastly, JARs secreted up to ∼181 ng IL-4 and ∼43 ng IL-13 following treatment with 600 ng mRNA. This data demonstrates that our LNP platform can induce secretion of cytokine mRNA in both 1^st^- and 3^rd^-trimester cell lines, and that each cell line secretes variable amounts of cytokine at the same administered doses, likely due to differences in the metabolic rate of these cell lines in culture.

**Figure 2.**
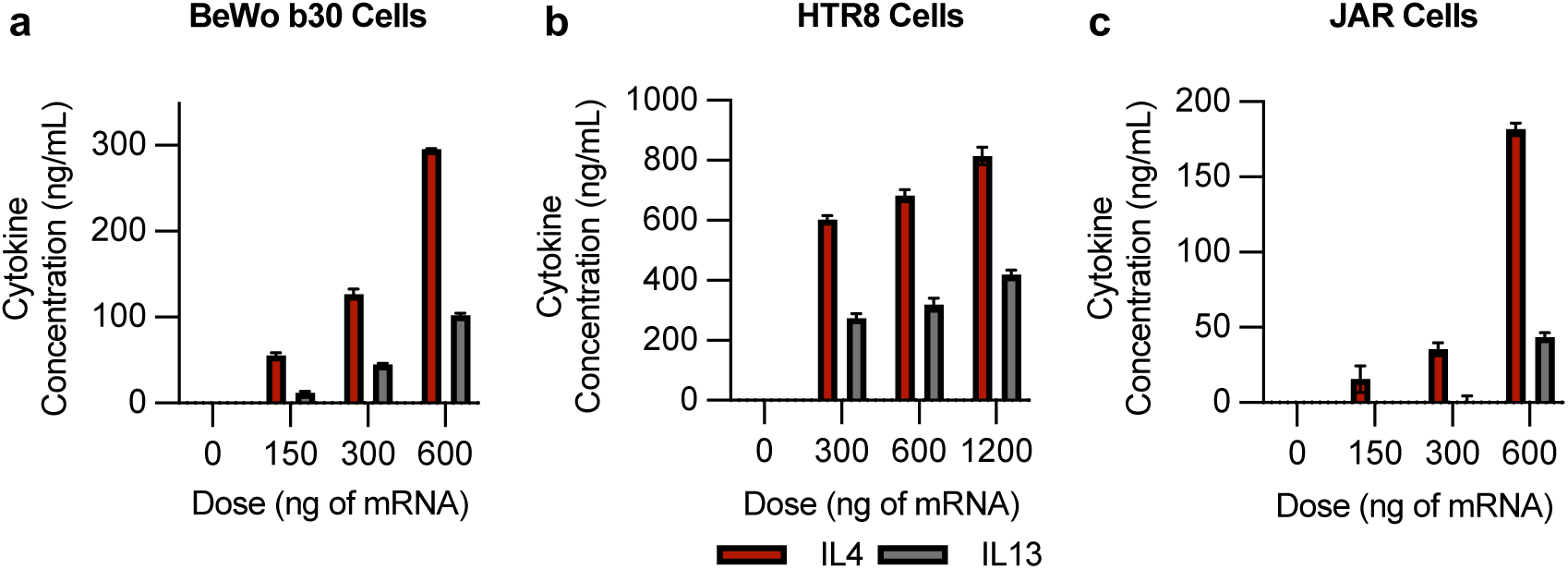
Ectopic cytokine production from trophoblast cell lines following LNP-mediated mRNA delivery. IL-4-LNPs and IL-13-LNPs elicit dose-dependent ectopic cytokine production in (a) BeWo b30 cells, (b) HTR8/svNEO cells, and (c) JAR cells. This CM was used to induce macrophage polarization, as assessed in later experiments.

We also evaluated cellular metabolic activity following LNP treatment as a measure of viability. We treated cells with 300 ng mRNA/well, which resulted in statistically insignificant changes in metabolic activity, demonstrating that the required LNP dose is nontoxic to cells (Supplementary Fig. 1). Importantly, this mRNA dose yields concentrations of secreted cytokines that exceed the amount required for polarization experiments, as described below. From this data, we determined that LNPs successfully delivered IL-4 and IL-13 mRNA to both BeWos and HTR8s with low toxicity at the required doses (<300 ng). BeWos had the largest dynamic range for both IL-4 and IL-13 secretion (Fig. 2). Further, the amounts of IL-4 and IL-13 secreted by BeWos treated with 150 ng mRNA aligns with the desired cytokine dose for macrophage polarization experiments.^56^ Thus, we used BeWos in the remaining experiments to investigate macrophage polarization.

### LNP-mediated cytokine secretion yields macrophage polarization *in vitro*

All of the experiments described herein were performed using BeWos. We treated BeWos in a T25 tissue culture flask with 2500 ng mRNA each of hIL-4-LNPs and hIL-13-LNPs (5000 ng mRNA total/flask). This dose is analogous to 100 ng/well based on our dose response experiments described above, but scaled up by cell number to generate sufficient levels of secreted cytokine for macrophage polarization (Fig. 3/Supplementary Figs. 2/3). This dose was chosen based on prior literature showing that 40 ng/mL of recombinant IL-4 and 20 ng/mL of recombinant IL-13 yield the highest expression of alternative activation markers on macrophages.^44,56,57^ BeWos were treated with PBS, hIL-4-LNPs, and/or hIL-13-LNPs and the conditioned media (CM) was collected after 12 hrs and added to human-derived monocytes; the CM-treated macrophages were compared to the PBS negative control and a recombinant peptide positive control. Macrophage phenotype was assessed after 48 hrs by flow cytometry. We first gated for CD14^+^CD11b^-^ macrophages and used this population of cells to analyze macrophage marker expression (Supplementary Fig. 2).

**Figure 3.**
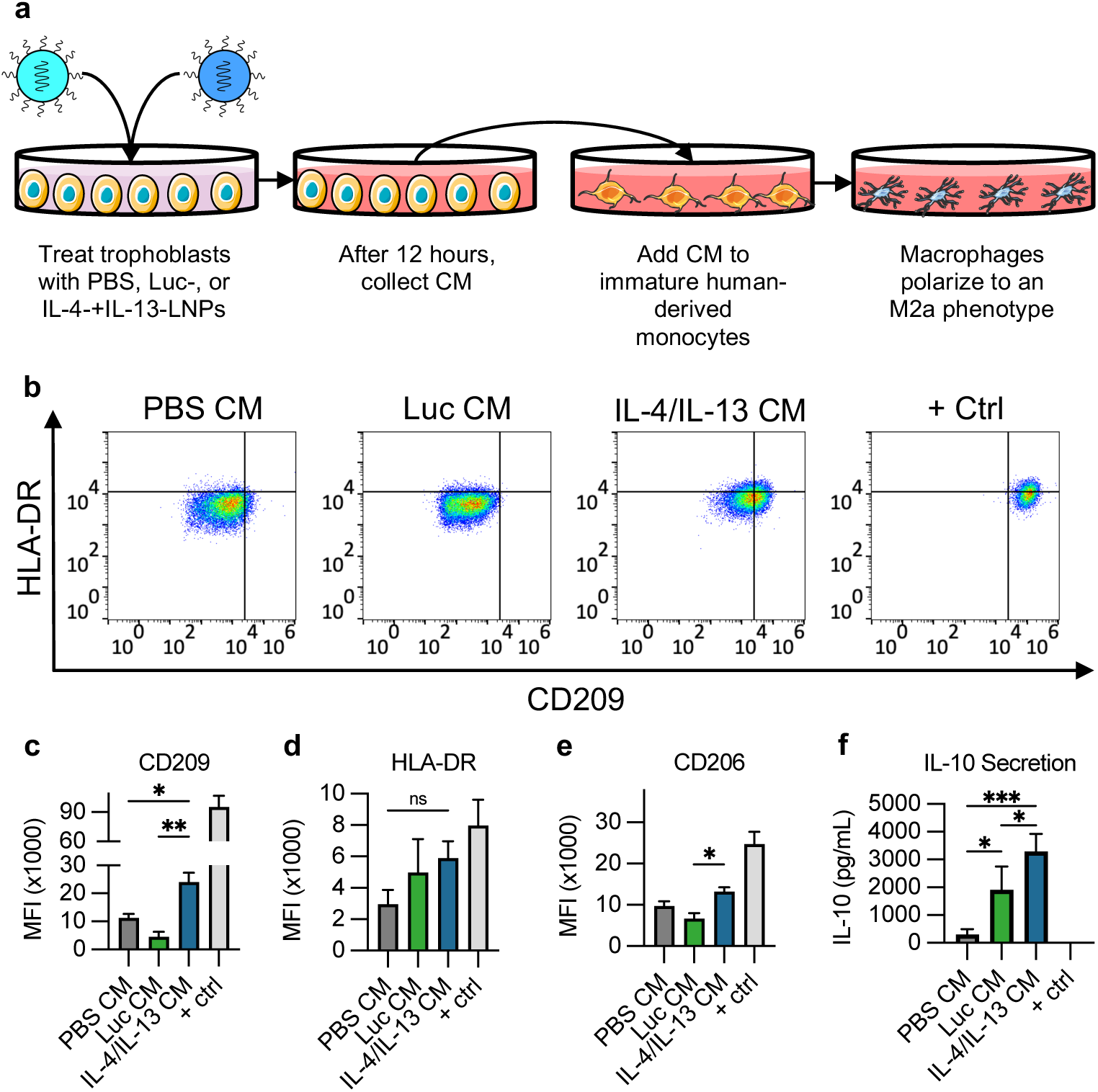
CM collected from BeWo cells treated with IL-4-LNPs and IL-13-LNPs induces polarization of patient-derived human macrophages. (a) Trophoblast cell lines were treated with IL-4-LNPs or IL-13-LNPs, and the conditioned media (CM) was assessed for cytokine content after 12 hours. (b) Representative flow cytometry data showing the phenotypic shift of patient-derived CD14+CD11c-monocytes after treatment with CM from treated BeWo cells. Flow cytometry analysis was used to assess (c) CD209 and (d) HLA-DR expression on CD14+CD11c-cells, and (e) CD206 expression on CD14+CD209+HLA-DR+ cells. CM-treated macrophages were assessed for (f) IL-10 secretion by ELISA. *p<0.05 and **p<0.002.

Macrophages cultured in CM from hIL-4-LNP and hIL-13-LNP-treated BeWo’s exhibited a marked upregulation of the M2 marker, CD209 (DC-SIGN), compared to CM from PBS- or Luc-LNP treated BeWos (Fig. 3b,c/Supplementary Fig. 3). There was no significant change in HLA-DR expression, which is often associated with pro-inflammatory macrophages, across any of the experimental groups (Fig. 3b,d/Supplementary Fig. 3).^58,59^ Further analysis of the CD14^+^CD11c^-^HLA-DR^-^ population showed increased expression of CD206 on macrophages treated with CM from BeWos treated with hIL-4-LNPs and hIL-13-LNPs compared to controls (Fig. 3e/Supplementary Fig. 3). CD206 is a mannose receptor expressed on the surface of anti-inflammatory macrophages.^60^ It has been shown in prior studies that macrophages with high levels of CD206 secrete more IL-10 than macrophages that lack CD206 expression, which made CD206 an interesting target receptor for our studies.^61^ These results demonstrate that BeWo CM can induce anti-inflammatory macrophage polarization approaching the positive control, in which monocytes were directly treated with IL-4 or IL-13 recombinant peptides (Fig. 3/Supplementary Fig. 3).

In addition to surface marker expression, we also assessed the ability for macrophages to secrete IL-10 after culture in the CM (Fig. 3f). IL-10 secretion is a hallmark of the alternatively activated macrophage phenotype.^45,62^ Macrophages that were cultured in CM from PBS-treated BeWos elicited low levels of IL-10 secretion (303.06±192.92 pg/mL). Macrophages that were cultured in CM from BeWos treated with Luc-LNPs secreted higher amounts of IL-10 (1513.3±340.8 pg/mL) compared to PBS-treated BeWos. We hypothesize that this increase in IL-10 was from LNPs activating other biological mechanisms that induce IL-10 secretion. For example, drug delivery systems, including the Pfizer-BioNTech mRNA vaccine and tumor-derived exosomes, have been shown to activate the STAT pathway and increase IL-10 secretion.^62–64^ Comparatively, macrophages that were cultured in CM from hIL-4-LNP and hIL-13-LNP treated BeWos elicited significantly increased IL-10 secretion (3453<u>±</u>808.6 pg/mL), more than double that of PBS or Luc-LNP-treated cells. Interestingly, macrophages exposed to recombinant peptides did not secrete any detectable IL-10 (Fig. 3f), which is likely due to the CM from LNP-treated BeWos eliciting a more robust anti-inflammatory response compared to the peptides.^41,65^ This data indicates that LNP-mediated cytokine secretion from trophoblasts can induce functional macrophage polarization, as assessed by IL-10 secretion. Further, LNP-mediated IL-10 secretion exceeds the positive control, demonstrating that LNPs can serve as a tool to modulate macrophage phenotypes *in vitro*.

### Macrophage polarization increases with decreasing dose of mRNA

While the treatment doses used above yielded significant macrophage polarization and IL-10 secretion, we aimed to increase the macrophage polarization status to more closely approach the positive controls treated with recombinant peptides. We hypothesized that there is an inverse relationship between cytokine dose and extent of polarization, such that lowering the doses of cytokines in the CM would increase the expression of markers of alternatively activated macrophages. This is because post-translational modifications have a significant impact on the potency and efficacy of protein-based drugs.^66,67^ With increased potency compared to the recombinant peptides, the original high doses of mRNA could lead to receptor saturation and internalization, or negative feedback loops, that preclude the extent of polarization.^68–71^

To test this hypothesis, we treated BeWos with a range of mRNA doses in LNPs to generate CM containing variable cytokine concentrations, allowing us to assess how cytokine concentration drives macrophage polarization. Based on the doses tested in our dose response study (Fig. 2), LNP-treated BeWos produced CM containing ∼127 ng/mL IL-4 and ∼45 ng/mL IL-13, without causing toxicity (Fig. 2/Supplementary Fig. 1), at the 300 ng mRNA dose. These cytokine concentrations are 3.2-fold and 2.2-fold higher, respectively, than the doses used in our initial polarization experiments (40 ng/mL IL-4 and 20 ng/mL IL-13). Since LNPs provided the ability to secrete excess of these cytokines from trophoblasts, we were able to evaluate a wide range of mRNA doses to elucidate the impact of cytokine dose on the extent of macrophage polarization.

LNPs were added to BeWos (0.5–0.03 ng mRNA/μL), the CM was added to human-derived monocytes, and surface expression of CD209 and CD206 was assessed. Our results show that decreasing the dose of mRNA in LNPs yields significantly increased surface expression of CD209 and CD206 in CD14^+^CD11b^-^HLA-DR^-^ macrophages (Fig. 4). At low mRNA doses of 0.06 ng/μL or 0.03 ng/μL, surface expression of CD209 or CD206, respectively, were statistically insignificant compared to the positive control peptides (Fig. 4). This confirms our hypothesis that there is an inverse relationship between administered cytokine dose and extent of polarization. Further, this data suggests that LNPs can be used as tools to induce ectopic secretion of cytokines to modulate macrophage phenotype, and that our approach is more potent than using recombinant peptides. This is beneficial towards clinical translation, as even low LNP delivery to the desired tissue could yield sufficient cytokine concentrations to impact macrophage activity.

**Figure 4:**
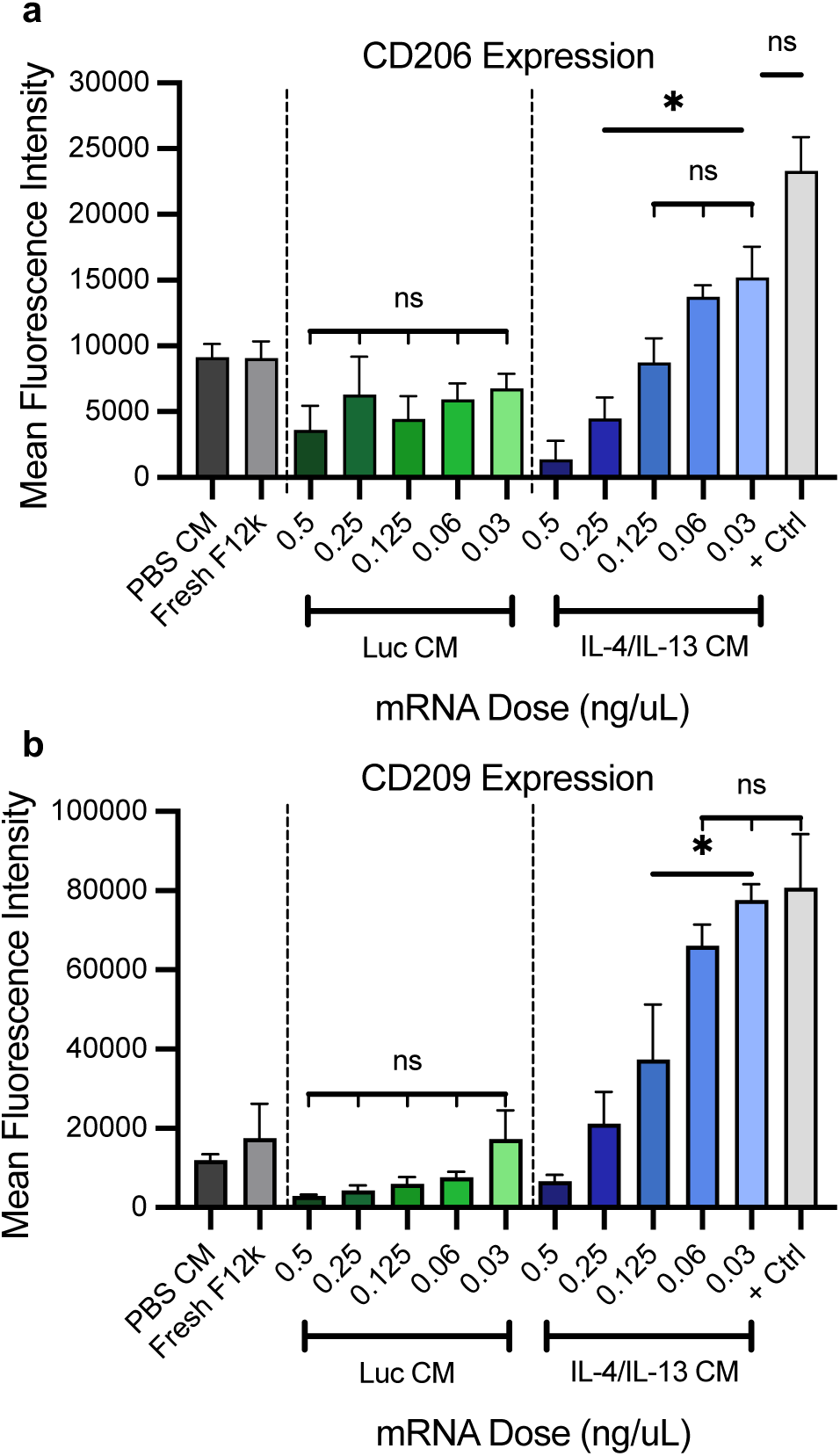
Decreasing the administered dose of mRNA leads to increased CD206 and CD209 expression. Cells were treated with IL-4-LNPs and IL-13-LNPs at doses ranging from 0.03–0.5 ng mRNA/µL. Flow cytometry was used to assess (a) CD206 expression on CD14+CD209+HLA-DR+ cells, and (b) CD209 expression on CD14+CD11c-cells. *p<0.05.

### *In vivo* LNP biodistribution in pregnant mice

In prior work, we demonstrated that the LNP formulation used here achieved placental delivery in late mouse pregnancy (gestational day (E) 17.5).^1^ Here, we aimed to assess placental delivery earlier in mouse gestation at E12.5. We are interested in evaluating LNP delivery at E12.5 because this is the earliest timepoint during murine gestation with a mature placenta and sufficient tissue volume for analysis.^72^ Pregnant female CD-1 mice were intravenously injected via the tail vein (0.5 mg/kg) with Luc-LNPs or PBS at E12.5. After 4 hr, mice were imaged using an *in vivo* imaging system (IVIS) (Fig. 5). Whole mouse images showed luciferase activity throughout the abdomen (Supplementary Fig. 4). Mice were euthanized and individual organs were imaged by IVIS. As expected, the liver and spleen had high luciferase expression (Fig. 5a). The LNPs induced luciferase expression in the placentae at this earlier timepoint without delivery to the fetuses, similar to our prior study (Fig. 5b,c/Supplementary Fig. 5).^1^ The lack of fetal delivery indicates that Luc-LNPs did not cross the placental barrier into the fetus. This data demonstrates that this LNP platform can be used to deliver mRNA to the murine placenta as early as E12.5.

**Figure 5.**
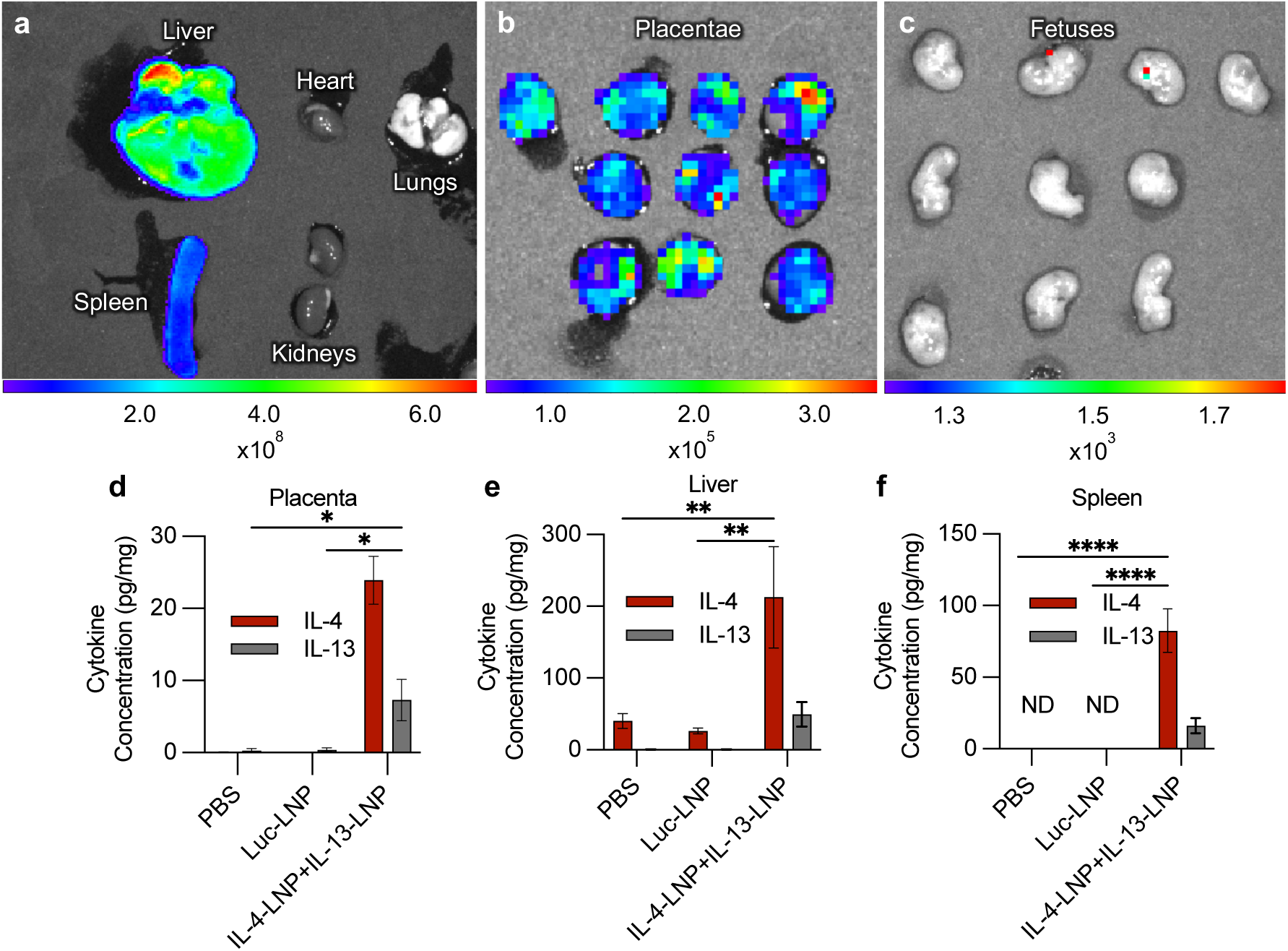
LNP delivery increases cytokine levels within the tissues of interest. (a-c) Representative IVIS images of (a) maternal tissues, (b) placentae, and (c) fetuses collected from dams 4 hours after Luc-LNP injections (0.5 mg mRNA/kg). (d-f) IL-4 and IL-13 levels in the (d) placentae, (e) liver, and (f) spleen 4 hours after injections of IL-4-LNPs+IL-13-LNPs (0.5 mg mRNA/kg). ND refers to raw data values below the limit of detection. *p<0.05, **p<0.01, and ****p<0.0001.

### mIL-4-LNPs and mIL-13-LNPs induce cytokine secretion in mouse placentas

To assess cytokine mRNA delivery, healthy pregnant CD-1 mice were intravenously injected with mIL-4-LNPs and mIL-13-LNPs, or PBS at E12.5. In these experiments, the mRNA encoded the mouse sequences of these cytokines, compared to the human mRNA used in the in vitro experiments described above. The livers, spleens, and placentas were collected 4 hrs post-injection and assessed for IL-4 and IL-13 cytokine concentrations (Fig. d-f). Overall, IL-13 secretion was lower than IL-4 secretion in all tissues, consistent with our *in vitro* results (Fig. 5d-f/Supplementary Fig. 6). Placentas from dams treated with PBS and Luc-LNPs had no detectable IL-4 or IL-13 expression, indicating that any amount of these cytokines in the placenta tissue is a result of LNP-mediated delivery. There was ∼24 pg/mg of IL-4 and ∼7 pg/mL of IL-13 in the placentae of IL-4-LNP+IL-13-LNP treated dams (Fig. 5d). Both livers and spleens had increased IL-4 (186 pg/mg and 82.5 pg/mg, respectively) and IL-13 (49 pg/mg and 16 pg/mg, respectively) compared to Luc-LNP treated dams (Fig. 5e,f).^1^ This data demonstrates that LNPs reach the placenta in sufficient levels at E12.5 to induce cytokine secretion. In future studies, we aim to further enhance placenta-specific delivery to promote cytokine secretion in the placenta while minimizing hepatic delivery.

### Assessing cytokine levels in mouse serum

To evaluate the immunomodulatory and immunotoxicity effects of the LNPs, we measured expression levels of various cytokines and chemokines in the serum 4- and 48- hr post-injection by Luminex. Notably, IL-6, IL-1a, and TNF-α did not change between treatment groups at either timepoint, suggesting that the LNPs did not elicit significant systemic pro-inflammatory responses (Fig. 6a,b Supplementary Figs. 6,7). In contrast, our data revealed increased IL-4, IL-13 (Fig. 6c), and MCP-1 4-hrs post-injection (p=0.134), and increased IL-4, IL-13, IL-10 (p=0.184) and MCP-1 (p=0.125) 48-hrs post-injection in dams treated with mIL-4-LNPs and mIL-13-LNPs compared to controls (Fig. 6a-d). MCP-1 is a potent chemoattractant responsible for recruiting monocytes to sites of tissue remodeling or inflammation.^73^ Alternatively, IL-10 is an anti-inflammatory cytokine crucial for resolving inflammation and promoting tissue repair.^29^ The elevation of IL-10 at 48-hr post-injection (p=0.184) aligns with our *in vitro* data showing that the LNPs induce IL-10 secretion (Fig. 3f), and is consistent with the expected effects of delivering IL-4 and IL-13 mRNA. Together, these results indicate that LNPs effectively promoted expression of chemokines important for immune cell recruitment, as well as the induction of IL-10, supporting the anti-inflammatory and tissue-repairing functions of the delivered IL-4 and IL-13 mRNA.

**Figure 6.**
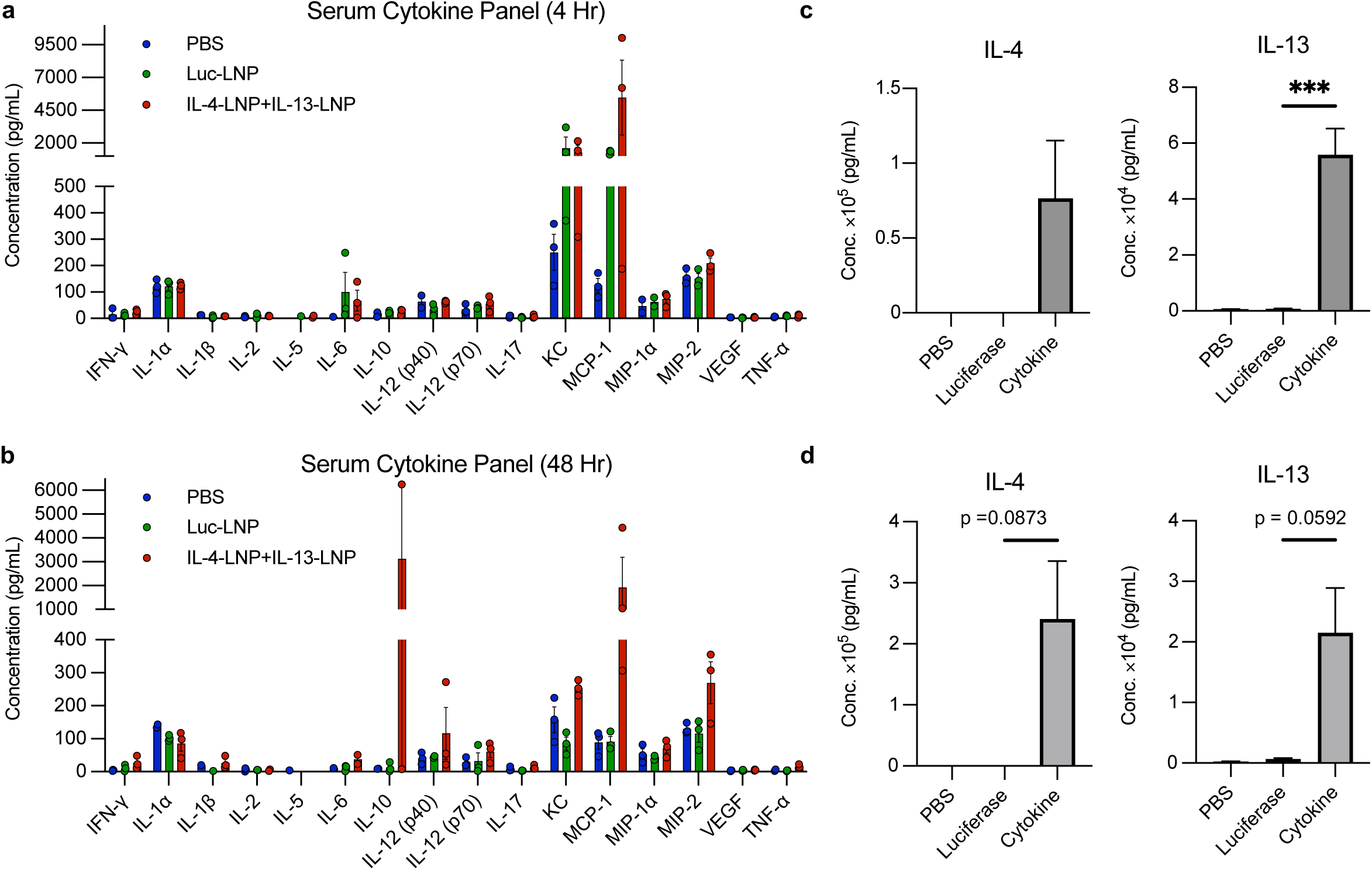
Cytokine levels in dam serum following LNP injections. (a,b) Luminex assay quantification of cytokine levels in dam serum (a) 4 hours and (b) 48 hours after injections of PBS, Luc-LNPs, or IL-4-LNPs+IL-13-LNPs. (c, d) IL-4 (left) and IL-13 (right) levels in the serum (c) 4 hours and (d) 48 hours after injections. *p<0.05, **p<0.01, ***p<0.001 and ****p<0.0001.

## Discussion

The development of therapeutics for use during pregnancy has only recently begun, but is an urgent unmet clinical need.^1,4,6–11^ The need for therapeutics to treat diseases of pregnancy is exemplified by recent industry attention, as Comanche Biopharma has developed an sFLT-1 siRNA, without a nanoparticle carrier, that is currently in clinical trials and received Fast Track designation from the FDA in August 2023.^74^ Our lab, and others, are interested in promoting placenta-specific nucleic acid delivery through the development of nanoparticle-based delivery strategies.^1,4,8,9^ Of note, prior studies have developed LNPs for delivering mRNA encoding angiogenesis-promoting factors as potential therapeutic agents for preeclampsia.^1,4,8^ For example, our lab has explored LNPs to deliver PlGF mRNA to the placenta to reverse the elevated sFlt-1:PlGF ratio characteristic in preeclamptic pregnancies.^1^ While prior studies have focused on gene therapies that aim to reverse preeclampsia symptoms, the current study represents a new approach that utilizes LNPs to induce ectopic cytokine secretion for immunomodulation in the placenta. Using LNPs to perturb immunological axes within the placenta offers a means to study and treat placental disorders before significant damage occurs.

Towards this goal, we used an LNP platform previously developed by our lab^1^ to deliver cytokine mRNA to tissue-resident cells, leading to local cytokine secretion. The importance of achieving controlled immunomodulation in the placenta is supported by studies suggesting that an immune mismatch at the maternal-fetal interface early in pregnancy is associated with abnormal placental development, including the poor spiral artery invasion characteristic of preeclampsia.^75^ In particular, there is decreased IL-4 and IL-10 in preeclamptic placentas and in the decidua of patients with recurrent pregnancy loss compared to healthy pregnancies, demonstrating their importance during pregnancy establishment.^76–78^ Our data demonstrated that LNPs can deliver IL-4 and IL-13 mRNA to trophoblasts, leading to cytokine secretion and subsequent polizarization of primary human monocytes towards anti-inflammatory macrophages *in vitro*. Our *in vivo* experiments show mRNA delivery to and cytokine secretion in mouse placentae. Further, there was increased IL-10 levels in the serum of mice treated with IL-4- and IL-13-LNPs, indicating that IL-4 and IL-13 secretion is able to induce this anti-inflammatory pathway. However, it is important to note that these experiments did not yield macrophage polarization within the placentas at the doses administered. Thus, there is much opportunity to iterate on our LNP design and dosing regimens to reduce serum levels and enhance placenta-specific delivery, towards the goal of achieving tissue-resident macrophage polarization.

Our data demonstrates that lower cytokine mRNA doses yielded significantly higher markers of macrophage polarization *in vitro* compared to our original treatment dose, which was based on prior literature.^44,56,57^ We hypothesize that this inverse relationship between cytokine dose and extent of polarization is due to post-translational modifications achieved by delivering the mRNA opposed to the recombinant peptides.^66,67^ Thus, delivering mRNA may yield more potent cytokines compared to recombinant peptides. High doses of the more bioactive molecules may lead to receptor saturation and internalization or negative feedback mechanisms that limit further macrophage polarization.^68–71^ This is supported by the fact that IL-4 binds to the IL-4 receptor with high affinity; thus, a relatively small amount of cytokine is necessary to occupy all surface receptors and activate downstream signaling.^79^ The ability to use LNPs to achieve robust immunomodulation at lower cytokine concentrations is critical towards the clinical translation of this technology because it could overcome some of the inherent type 1 inflammation associated with LNP therapies.^80,81^ Further, lower dosing regiments would decrease the potential for adverse effects to both maternal and fetal tissues.

Here, we present the first application of LNPs to induce cytokine secretion in the placentae of mice, demonstrating their potential to be used as tools to study the local effects of cytokines within the tissue of interest. Our approach to use LNPs to deliver cytokine mRNA to tissues, such as the placenta, can overcome challenges associated with peptide or protein delivery. Cytokines have been used as therapeutic agents for the treatment of many diseases, most notably cancer, infectious disease, and autoimmune diseases.^82^ The *in vivo* efficacy and clinical translation of cytokine therapies is limited to short half-life and off target effects.^82–84^ Using LNPs to deliver cytokine mRNA, rather than recombinant peptides, offers an alternative approach for local cytokine therapy for diseases of pregnancy and for other diseases. By enabling local immunomodulation, this approach could provide new approaches and enable foundational understanding of the role of immune activity in specific tissues and across a myriad of diseases, such as the placenta.

## Materials and Methods

### LNP Formulation

The C12-200 ionizable lipid was fabricated in house.^85,86^ Other LNP components including cholesterol, DOPE, and DMPE-PEG2000 (ammonium salt)) were purchased from Avanti Polar Lipids Inc. (Birmingham, AL). Codon optimized mRNA was prepared by in vitro transcription through a collaboration with the Engineered mRNA and Targeted Nanomedicine core facility at the University of Pennsylvania (Philadelphia, PA).

Human IL-4 (hIL-4), human IL-13 (hIL-13), mouse IL-4 (mIL-4), and mouse IL-13 (mIL-13) were synthesized with 1-methylpseudouridine modifications, co-transcriptionally capped using the CleanCap system (TriLink), and purified using cellulose based chromatography.^87^ LNPs were formulated by combining one volume of an organic phase containing the lipid components in ethanol and three volumes of the aqueous phase containing mRNA by rapid mixing with micropipettes. mRNA was diluted in citrate buffer (pH 3) to an mRNA:ionizable lipid weight ratio of 1:10 for all LNP formulations. The LNPs consisted of 53.5% cholesterol, 35% C12-200, 10% DOPE, and 1.5% DSPE-PEG-2000 by molar weight ratio. After mixing, the LNPs were dialyzed against PBS (pH 7.2) for 1 hr, sterile filtered using 0.2 µm filters, and stored at 4°C until use.

### LNP Characterization

Dynamic light scattering (DLS) measurements and mRNA encapsulation efficiency were assessed as previously described.^1^ The average and standard deviation of the hydrodynamic diameter, polydispersity index, and encapsulation efficiency of each LNP is reported. For DLS, each LNP formulation was diluted 1:100 in deionized water in cuvettes and measurements were run on a Malvern Zetasizer Nano ZS (Malvern Panalytical). Encapsulation efficiency was calculated using the QuantiFluor® RNA System (Promega). Briefly, LNPs were diluted 1:100 in 1X Tris-EDTA (TE) buffer in two microcentrifuge tubes for each LNP formulation. 1% v/v Triton X-100 (Thermo Scientific) was added to one tube and both were heated to 37°C and shaken at 600 rpm for 1 min, followed by cooling to room temperature for 10 mins. LNP samples and RNA standards were plated in triplicate in black 96-well plates, the fluorescent reagent was added, and intensity was measured using a plate reader (Molecular Devices) per manufacturer instructions (excitation, 492 nm; emission, 540 nm). Background signal was subtracted from each well and triplicate wells for each LNP were averaged. RNA content was quantified by comparing samples to the standard curve, and encapsulation efficiency (%) was calculated according to the equation below, where A is the RNA content in samples without Triton X-100 (intact LNPs) and B is the RNA content in samples with Triton X-100 (lysed LNPs).

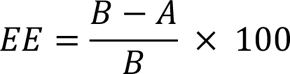

### Cell Culture

BeWo b30 subclone (BeWo) cells were cultured in Kaighn’s F-12k with L-glutamine (F12k) media (Corning) supplemented with 10% fetal bovine serum (FBS, Avantor) and 1% penicillin/streptomycin (pen/strep, VWR). HTR8/svNEO cells were cultured in RPMI-1640 (Cytiva) supplemented with 10% fetal bovine serum (FBS, Avantor) and 1% penicillin/streptomycin (pen/strep, VWR). HTR8/svNEO and BeWos were cultured in either 24-well plates or 25 cm^2^ (T25) flasks at plating densities of 1.2×10^5^ cells/well or 1.0×10^6^ cells/flask, respectively. Peripheral human monocytes collected from healthy donors were purchased from The Human Immunology Core at the University of Pennsylvania and cultured in RPMI-1640 media supplemented with 10% FBS and 1% pen/strep in 24-well plates. Cultures were grown in an incubator set at 37 °C with 5% CO_2_.

### LNP Delivery to Trophoblasts *in vitro*

BeWo, HTR8/svNEO, and JAR cells were plated at 1.2 x 10^5^ cells/well in 24-well plates and incubated for 6 hr to adhere. After 6 hr, IL-4-LNPs or IL-13-LNPs were added into each well using a micropipette and gently mixed by lateral translation of the plate. Treated cells were incubated for 24 hr, after which media containing secreted cytokines was collected and stored in microcentrifuge tubes (Thomas Scientific) at −80 °C for quantification by ELISA. Cytokine content in samples was assessed using IL-4, IL-13, and IL-10 ELISA assays (ThermoFisher). Samples were thawed on ice, mixed, and diluted in ELISA assay buffer. Appropriate dilutions were determined by serially diluting samples 0-1000-fold in assay buffer. Samples of cytokine-containing media were prepared following manufacturer instructions for each ELISA kit and absorbance was measured using a plate reader at 450 nm. Absorbance was compared to a standard curve to quantify the amount of cytokine in each sample. Statistical analysis was performed using GraphPad to assess significance by one-way ANOVAs with posthoc Tukey (*p<0.05, **p<0.01, ***p<0.001, ****p<0.0001).

### Macrophage Polarization

Human monocytes were cultured at 2.0×10^6^ cells/well in a 24-well plate in RPMI-1640 supplemented with macrophage colony stimulating factor (M-CSF), FBS, and pen/strep. Media was exchanged with fresh media after 72 hr, and the cells were cultured for a total of 5 days prior to the experiments described below. Concurrently, BeWos were seeded at 1.0×10^6^ cells/flask in T25 flasks in F12k media and allowed to adhere for 8 hr. After 8 hr, BeWos were treated with PBS, Luc-LNPs, or IL-4-LNPs and IL-13-LNPs at a dose of 0.5 ng/μL total mRNA (0.25 ng/μL each) for 12 hr. Next, 1.0 mL of conditioned media from LNP-treated BeWos, or media containing IL-4 (40 ng/ml) or IL-13 (20 ng/ml) recombinant peptides as controls, was transferred onto the monocytes. Unless otherwise noted, all treatment groups included supplementation with M-CSF (40 ng/mL) (PeproTech) throughout the experiments. Lastly, macrophage polarization was assessed by flow cytometry, as explained below. Statistical analysis was performed using GraphPad to assess significance by one-way ANOVAs with posthoc Tukey (*p<0.05, **p<0.01, ***p<0.001, ****p<0.0001).

### *In vivo* Biodistribution Studies in Dams

Female CD-1 mice between 8-39 weeks of age were maintained, bred, and used in accordance with Animal Use Protocols approved by the Institutional Animal Care and Use Committee at the University of Delaware (AUP #1320 and #1341). Mice at gestational day (E) 12.5 were intravenously injected via the tail vein with Luc-LNPs, suspended in sterile PBS, at a dose of 0.5 mg mRNA/kg mouse weight. After 4 hr, dams were anesthetized via isoflurane and injected intraperitoneally with d-luciferin potassium salt (150 mg/kg) (Biotium). Anesthetized dams were placed supine into the *in vivo* imaging system (IVIS) (Lumina III, PerkinElmer). After whole body imaging, dams were euthanized and the placentae, fetus, liver, lung, uterus, heart, and spleen were collected and imaged. Raw luminescence (radiance in photons/sec/cm^2^/sr) is reported without normalization.

### IL-4-LNP and IL-13-LNP Delivery *in vivo*

Female CD-1 mice at E12.5 were intravenously injected via the tail vein with 0.5 mg mRNA/kg mouse weight of IL-4-LNPs and IL-13-LNPs, Luc-LNPs, or sterile PBS at an equivalent volume. After 4 or 48 hr, mice were euthanized and the placentae, fetus, liver, lung, uterus, heart, and spleen were collected. Whole blood was obtained by cardiac puncture and stored in tubes containing 0.5 M EDTA on ice until analysis by ELISA as described above. Tissues were immediately placed in RPMI-1640 supplemented with 10% FBS (Avantor) and 1% pen/strep (VWR) on ice. Following dissections, tissues were rinsed with PBS, cut using surgical scissors, and ground until paste-like in tissue grinding tubes (VWR). After mechanical digestion, enzyme digestion mix (5% FBS, 2mg/mL Collagenase A, and 28 U/mL DNase I) was added to each tube and shaken at 37°C for 45 min. The digested tissue was passed through a cell strainer with a mesh size of 100 μm, which was then rinsed with complete media (DMEM with 5% FBS and 1% pen/strep). The filtrate was collected and subsequently passed through a cell strainer with a mesh size of 40 μm. The cell-containing filtrate was centrifuged at 400 x g for 5 min, the supernatant was aspirated, and the cell pellet was resuspended in RBC lysis buffer and incubated at room temperature for 5 min. PBS was added to RBC lysis buffer solution to quench the reaction. Cells were then centrifuged at 400 x g for 5 min at 4°C and supernatant was discarded. Cells were rinsed 2X with FACS buffer before being fixed using 3.7% formaldehyde for 15 min. Formaldehyde was removed and cells were stored in FACS buffer or fluorescent antibody staining and analysis for polarization by flow cytometry, as described below.

### Flow Cytometry

Flow cytometry was used to assess macrophage polarization following *in vitro* and *in vivo* experiments. For *in vitro* experiments, media was removed from macrophages and cells were lifted using Accutase in DPBS (Innovative Cell Technologies). Cells were centrifuged at 300 x g for 5 min in 1.5 mL microcentrifuge tubes (Thomas Scientific). The Accutase was aspirated and cells were rinsed twice with sterile FACS buffer (Rockland) before being plated into 96-well FACS plates (Avantor). Cells were stained for CD14 (BV650), CD11c (PE-Cy7), CD206 (PE), CD209 (BV421), and HLA-DR (AlexaFluor 488) using fluorophore tagged primary antibodies (BD Biosciences) for 45 mins. For *in vivo* experiments, cells were stained for F4/80 (BV650), CD11c (PE-Cy7), CD209 (BV421), and CD86 (AlexaFluor 700) using fluorophore tagged primary antibodies (BD Biosciences) for 45 min. Following staining, cells were centrifuged at 500 x g for 5 minutes, supernatant was removed, and cells were washed twice in FACS buffer to remove unbound antibodies. Stained cells were analyzed on a SA3800 Spectral Analyzer (Sony) and data was analyzed using FlowJo software (BD Biosciences). Statistical analysis was performed using GraphPad with significance determined by one-way ANOVAs with posthoc Tukey (*p<0.05, **p<0.01, ***p<0.001, ****p<0.0001).

### Luminex Assay

A Luminex assay was used to measure protein levels in the serum of treated mice. Mice were euthanized via CO_2_ asphyxiation followed by cervical dislocation. Whole blood was collected by cardiac puncture. Serum was separated from the whole blood by spinning the whole blood at 1000 x g for 10 minutes. Serum was stored at −80 °C until assay was performed. Serum samples were sent to the Human Immunology Core at The University of Pennsylvania for processing. Samples were prepared according the to the protocol provided by the manufacturer. Briefly, samples were diluted 2-fold or 20-fold in assay diluent before beginning the assay. The assay was performed on a FLEXMAP 3D luminex instrument by the staff of the human immunology core using their internal protocol. Statistical analysis was performed using GraphPad. One-way ANOVA with posthoc Tukey * p < 0.05.

## Author Contributions

S.I.H. and R.S.R. conceived the project ideas, designed and led experimental procedures, and wrote the manuscript. S.I.H., L.A.F., R.E.Y., T.V., N.B., K.M.N., and R.S.R. performed the experiments and analyzed the data. M.G.A. and D.W. contributed experimental resources and provided input on experimental design. R.S.R. and J.P.G were responsible for project supervision and funding acquisition. All authors discussed the results and data analysis and edited the manuscript.

## Acknowledgements

This work was supported by the Peter Joseph Pappas Fund through the Preeclampsia Foundation, the New Jersey Health Foundation (PC 44-22), the New Jersey Department of Health (COCR22PRG012, COCR23PRF027), the National Science Foundation (NSF) Engineering Research Initiation Program (Award #2301919), the NSF Graduate Research Fellowship Program (Award #2018266781), and the National Institute of Health (U19AI158930).

