## Supplemental Files for "Cytokine mRNA Delivery and Local Immunomodulation in the Placenta using Lipid Nanoparticles"

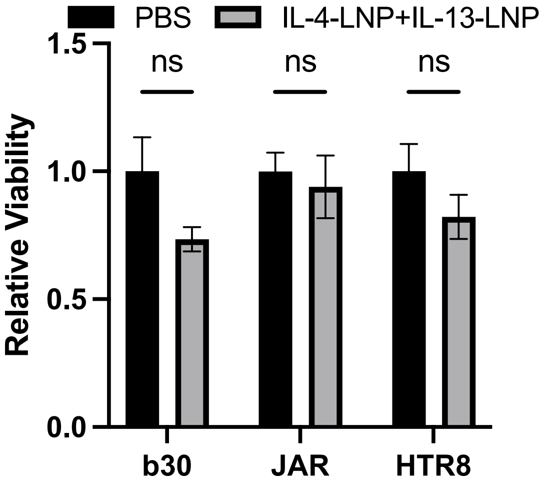


Supplemental Figure 1. Cell metabolic activity of each trophoblast cell line as assessed by MTS assay following treatment with IL4-LNPs and IL13-LNPs at 0.3 ng mRNA/μL. Data normalized to PBS treatment in each cell line.

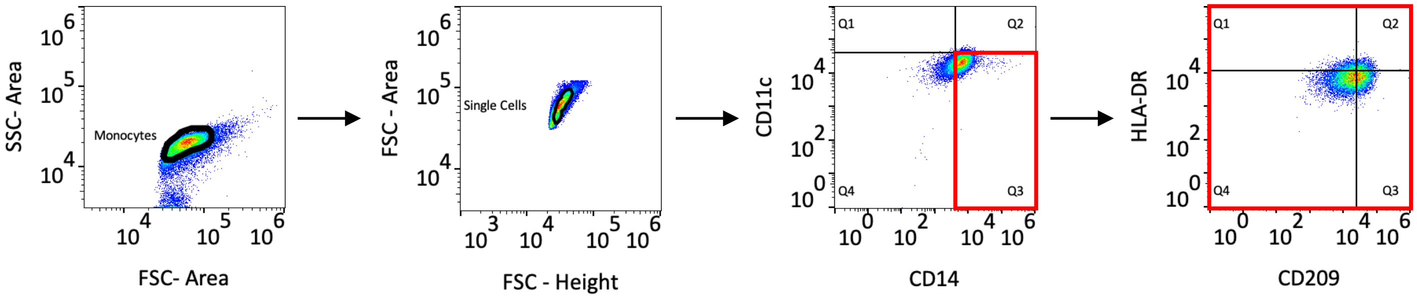


Supplemental Figure 2: Flow cytometry gating protocol used to quantify HLA-DR and CD209 on macrophages.

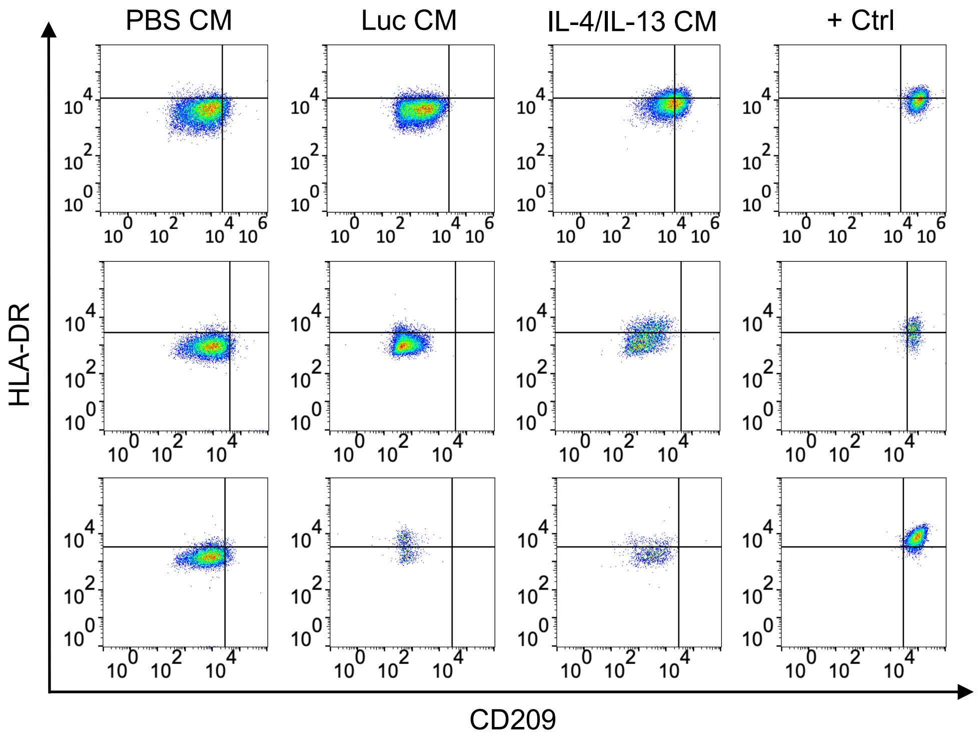


Supplemental Figure 3. Flow cytometry data from CD14+CD11c- populations of primary patient derived monocytes. Each row represents a biological replicate of monocytes collected from a different patient.

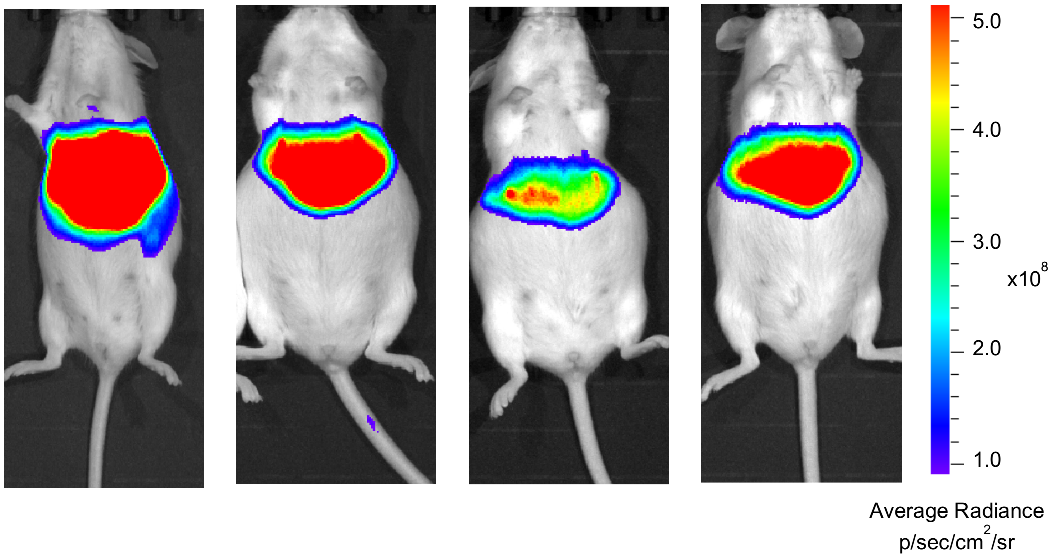


Supplemental Figure 4: IVIS images from each pregnant mouse injected with 0.5 mg/kg Luc-LNPs 4 hours after injection. Each column represents a biological replicate.

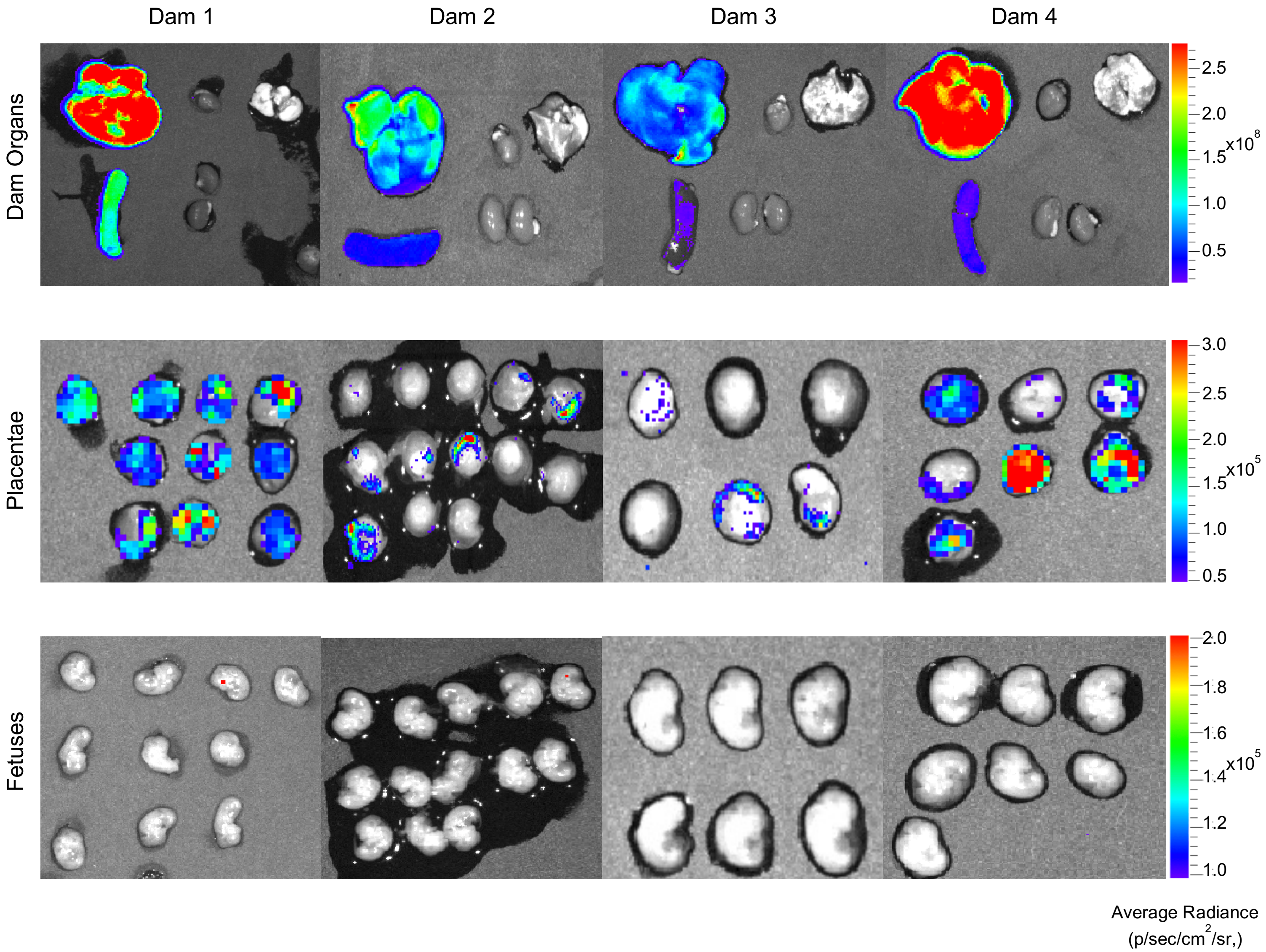


Supplemental Figure 5. IVIS images from each dam 4 hours after injection with 0.5 mg/kg Luc-LNPs. Each row represents a biological replicate.

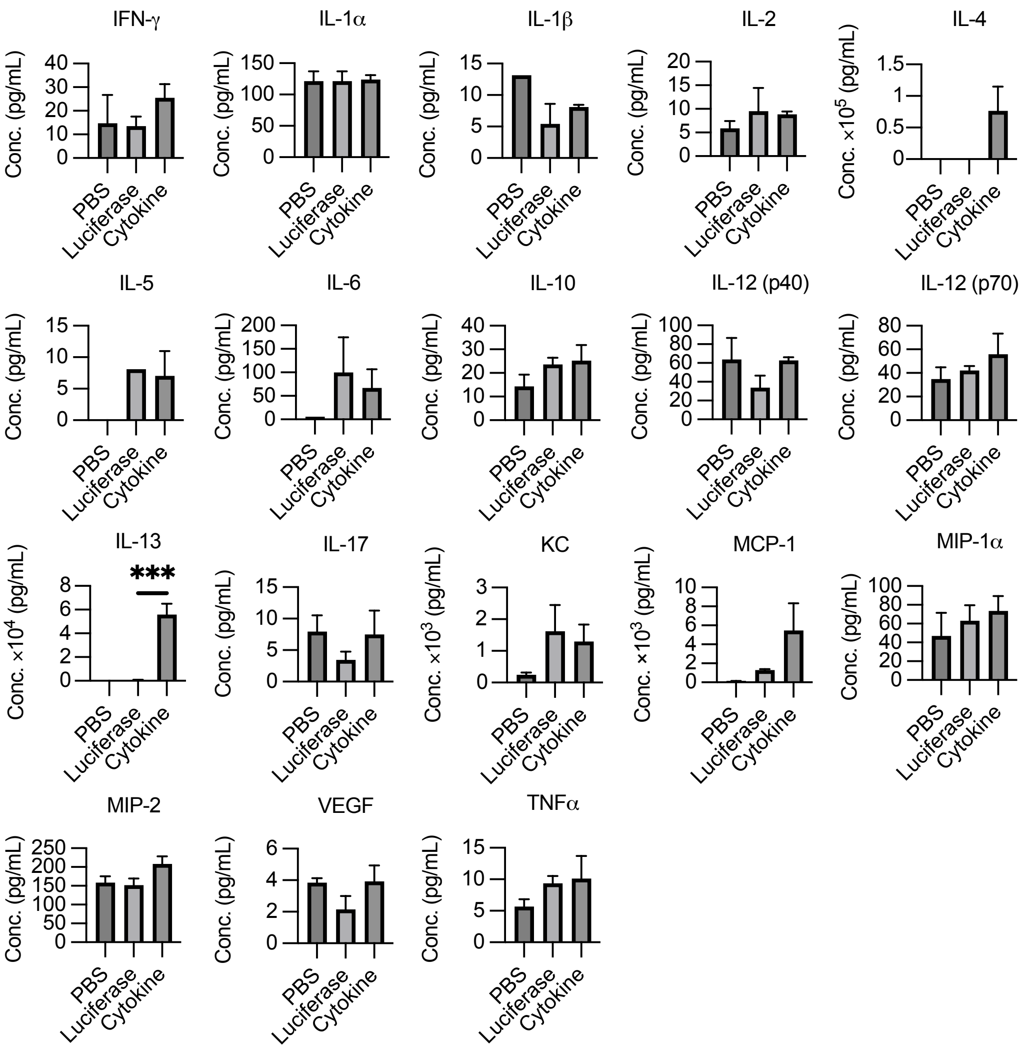


Supplemental Figure 6. Luminex or ELISA (IL-4 only) results showing cytokine levels in serum collected from dams 4 hours after LNP injection with individual one-way ANOVA performed on each serum analyte. ***p<0.001, ****p<0.0001.

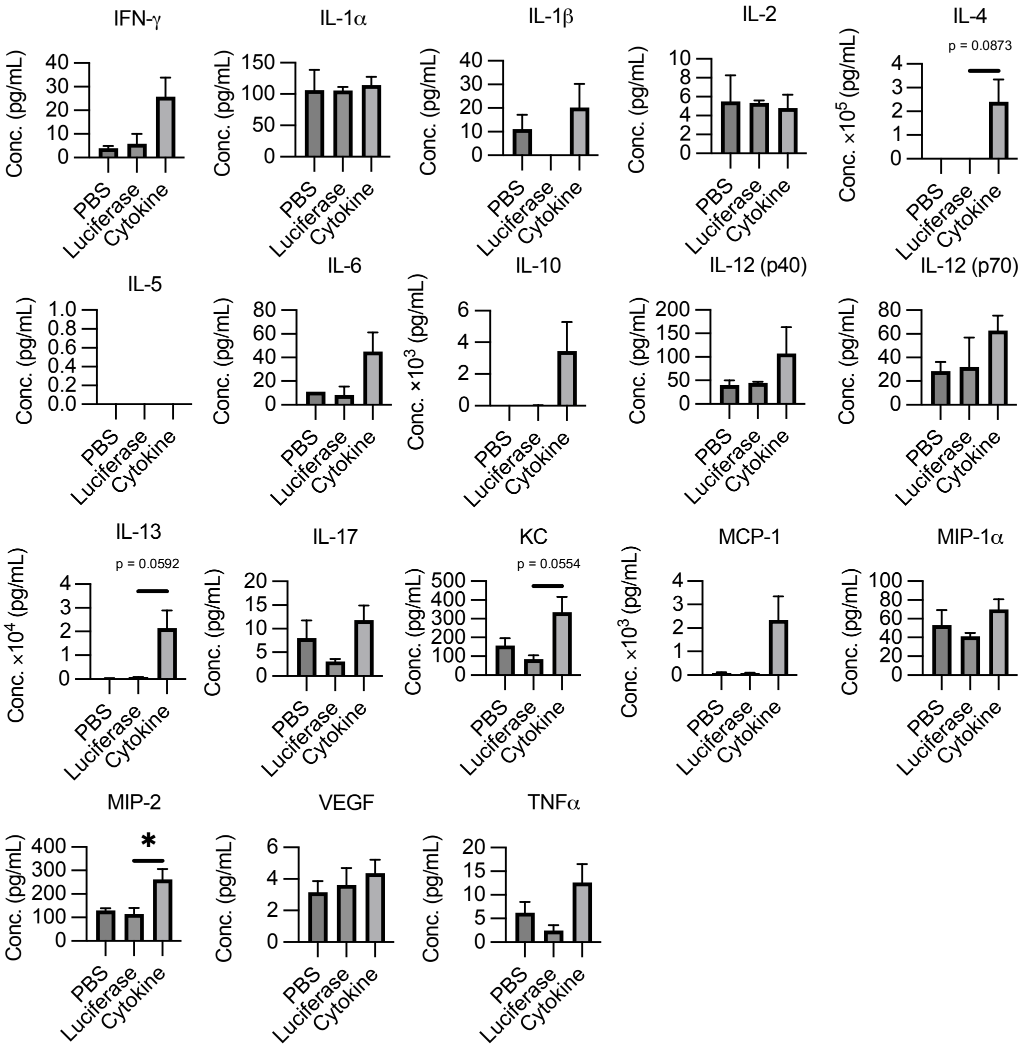


Supplemental Figure 7. Luminex or ELISA (IL-4 only) results showing cytokine levels in serum collected from days 48 hours after LNP injection with individual one-way ANOVAs performed on each serum analyte. *p<0.05.
